# Loss of neuronatin increases susceptibility of SERCA to thermal inactivation

**DOI:** 10.1101/2023.11.23.568317

**Authors:** Michael Barfoot, Val A. Fajardo, Jessica L. Braun

## Abstract

- Phospholamban and sarcolipin are two key regulators of the sarco(endo)plasmic reticulum Ca^2+^-ATPase (SERCA) that have been shown to protect SERCA from thermal inactivation in muscle
- We have recently detected neuronatin (NNAT) in murine skeletal muscle and have shown that it too regulates SERCA
- Here, we questioned whether NNAT would also protect SERCA from thermal inactivation
- In response to 60 min of heat stress at 40°C, maximal SERCA activity was significantly reduced in soleus homogenates obtained from NNAT knockout mice (-23%) but not wild-type mice (-5%).
- Our results add further support for NNAT’s role in regulating SERCA function in murine muscle

## Introduction

The sarco(endo)plasmic reticulum Ca^2+^-ATPase (SERCA) catalyzes the active transport of Ca^2+^ from the cytosol into the lumen of the sarco(endo)plasmic reticulum (SR/ER). In muscle, there are two main isoforms of SERCA: SERCA1a and SERCA2a (the fast and slow isoforms, respectively), with their functions being necessary for eliciting muscle relaxation, ensuring sufficient Ca^2+^ load for subsequent contractions, and maintaining low intracellular Ca^2+^ ([Ca^2+^]_i_) levels [1]. Thus, SERCA is critical for several physiological processes in muscle, and in many instances, SERCA dysfunction can lead to muscle wasting and weakness [2-4].

Structurally, the SERCA pumps contain vulnerable cysteine, lysine and tyrosine residues that are highly susceptible to free radical attack from various reactive oxygen/nitrogen species (RONS), which can alter protein structure and lower catalytic activity [5-7]. To preserve SERCA function, various chaperone and regulatory proteins can physically interact with SERCA, protecting it from RONS modification and impairment. These include heat shock protein 70 (HSP70) [8, 9], sarcolipin (SLN) and phospholamban (PLN) [10], all of which have been shown to protect SERCA against the ensuing rise in RONS associated with heat stress.

SLN and PLN are two small peptides in muscle that are known to regulate SERCA by reducing its affinity for Ca^2+^ - though SLN has the additional role of uncoupling SERCA-mediated Ca^2+^ transport from ATP hydrolysis (for review, see [11]). Recently, we have detected the presence of neuronatin (NNAT), which shares sequence homology with both SLN and PLN, in murine skeletal muscle where it binds to both SERCA1 and SERCA2 isoforms [11, 12]. Based on the sequence homology with SLN and PLN, we first posited that NNAT would regulate SERCA in a similar manner to these peptides; and our recent evidence provided in HEK cells indicates that like SLN, NNAT acts to uncouple the SERCA pump, potentially highlighting a role in muscle-based thermogenesis [12]. To extend our work and to further showcase NNAT as a SERCA regulatory protein in muscle, we questioned whether NNAT could protect SERCA from thermal inactivation similar to that previously found with PLN and SLN [10].

## Results and Discussion

To this end, we collected soleus muscles from male wild-type (WT) and NNAT knockout (NNAT^KO^) mice. This muscle was chosen as we have previously shown that NNAT is most abundant in the soleus compared to other skeletal muscles [11, 12]. The *Nnat* gene is paternally inherited and thus heterozygous *Nnat* deletion, with the paternal allele harboring the deleted gene, sufficiently knocks out NNAT protein in a variety of tissues [13]. Here, we show that this also holds true in murine skeletal muscle, as paternal heterozygous deletion of *Nnat* drastically reduced NNAT protein content (Figure 1A).

**Figure 1.**
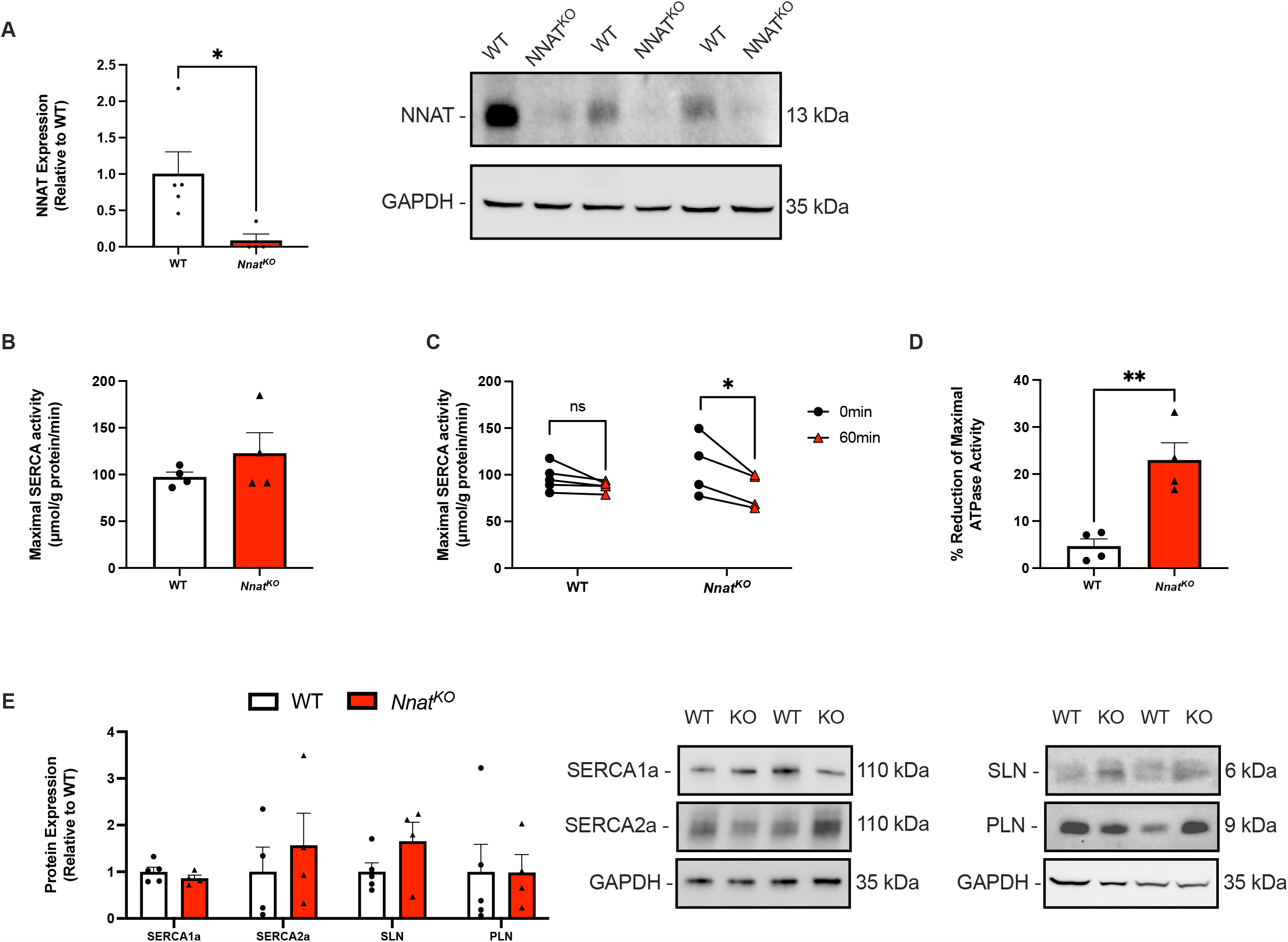
NNAT deletion accelerates thermal inactivation of SERCA in murine soleus muscle homogenates. **A)** Western blot analysis of NNAT protein content in soleus muscles obtained from wild-type (WT) and NNAT heterozygous knockout mice (NNAT^KO^) with the paternal allele harboring the genetic deletion of *Nnat*. **B)** Baseline maximal SERCA activity in soleus muscle homogenates measured at a *p*Ca of 5.0 (10 μM free [Ca^2+^]_i_). **C)** Maximal SERCA activity at baseline and after 60 min of heat stress (40°C incubation). **D)** % reduction in maximal SERCA activity after 60 min of heat stress in soleus muscles from WT and NNAT^KO^ mice. **E)** Western blot analysis of SERCA1, SERCA2, SLN and PLN in soleus muscles from WT and NNAT^KO^ mice. \**P* < 0.05, *\*\*P*<0.01. For A and D, a Student’s t-test was used and for C, paired t-tests were used. Data are presented as means ± SEM.

We then examined maximal SERCA activity in the soleus homogenates using an enzyme-linked spectrophotometric assay [14, 15] at a *p*Ca of 5.0. In a basal state, there were no differences in SERCA V_max_ between WT and NNAT^KO^ mice (Figure 1B). However, after incubating the homogenates for 60 min at 40°C, we found that while SERCA V_max_ was mostly preserved in the WT soleus (-5% reduction that was not statistically significant), maximal SERCA activity was significantly reduced in the NNAT^KO^ soleus (-23%) (Figure 1C). Furthermore, comparing the changes in maximal SERCA activity between WT and NNAT^KO^ mice also yielded a statistically significant difference (Figure 1D). These results were observed without any changes in the protein levels of SERCA1, SERCA2, SLN or PLN (Figure 1E).

In summary, our results add further evidence in support of a role for NNAT in regulating SERCA in muscle. Future studies aimed at investigating the effects of NNAT deletion or overexpression in various models of muscle disuse, disease or exercise training are needed to further uncover the physiological role of NNAT in muscle.

## Methods

### Animals

We purchased commercially available NNAT^KO^ mice from the Mutant Mouse Resource and Research Center (0324932-UCD). After cryorecovery, breeders were sent to Brock University to establish a breeding colony. For this study, we used male wild-type (WT) and heterozygous NNAT^KO^ mice with the paternal gene harboring the genetic deletion of *Nnat*. Mice were maintained on a 12h:12h light:dark cycle and were fed standard chow and water ad libitum. All protocols were approved by Brock University’s Animal Care Committee (AUP# 21-04-04) in accordance with the Canadian Council of Animal Care.

### Tissue collection

Soleus muscles were collected from 3-6 month old adult mice after euthanizing them via cervical dislocation while under general anesthesia (vaporized isoflurance). After collection, soleus muscles were flash frozen in liquid N_2_ and stored at -80°C until further use.

### SERCA activity and acute heat stress

To assess maximal SERCA activity with and without heat stress soleus muscles were first homogenized in homogenizing buffer (250 mM sucrose, 5 mM HEPES, 0.2 mM PMSF, 0.2% [w/v] NaN_3_; pH 7.5). Next, maximal SERCA activity was measured using an enzyme-linked spectrophotometric kinetic assay at a *p*Ca of 5 as previously described [14]. We conducted the assay at baseline and after 60 min of acute heat stress (40°C incubation).

### Western blotting

Western blotting was done to measure the protein levels of NNAT, SERCA1, SERCA2, SLN and PLN in soleus muscle homogenates from WT and NNAT^KO^ mice as previously described [3]. For SERCA1 and SERCA2, standard glycine-based gel electrophoresis was applied with polyvinylidene difluoride (PVDF) and the following antibodies: SERCA1 (MA3-911; ThermoFisher Scientific), SERCA2 (MA3-919; ThermoFisher Scientific). For SLN, PLN and NNAT a tricine-based gel system was used along with PVDF (PLN and NNAT) or nitrocellulose (SLN) membranes and the following antibodies: PLN (MA3-922; ThermoFisher Scientific), SLN (ABT13; Sigma-Aldrich) and NNAT (78122S; Cell Signaling Technology). To image the membranes, we used Immobilon^®^□ECL Ultra Western HRP Substrate (WBKLS0500; Burlington, MA, USA) or ThermoFisher Supersignal Femto West (PI34096; Rockford, IL, USA) and a BioRad ChemiDoc Imager. GAPDH (60004-1; Proteintech) was used as a load control.

### Statistics

Comparisons between baseline and post 60 min of acute heat stress were made with a paired t-test within a specific genotype. Comparisons between genotypes were made with a Student’s t-test. Statistical significance was set to *P* < 0.05. All data presented are means ±SEM.

## Acknowledgements

This work was funded by an NSERC Discovery Grant awarded to VAF. MB is supported by an Ontario Graduate Scholarship. HLB is supported by a CIHR Doctoral Award. VAF is supported by a Canada Research Chair Tier II Award in Tissue Plasticity and Remodelling Throughout the Lifespan.

## Data Availability

Data can be shared by the corresponding author upon reasonable request.

